# Genetically encoded SpyTag enables modular AAV retargeting via SpyCatcher-fused ligands for targeted gene delivery

**DOI:** 10.1101/2025.08.22.671696

**Authors:** Anja Armbruster, Maximilian Hörner, Hanna J Wagner, Claudia Fink-Straube, Wilfried Weber

## Abstract

Recombinant adeno-associated viral (rAAV) vectors are a leading platform of *in vivo* gene therapy, valued for their excellent safety, broad serotype diversity, and scalable production. Targeted delivery through capsid display of ligands holds great promise, yet current retargeting strategies often rely on extensive capsid re-engineering and restrict the use of ligands incompatible with intracellular expression systems. Here, we present a modular AAV retargeting platform that, for the first time, employs the SpyTag/SpyCatcher system via genetic integration into the AAV2 capsid. SpyTag is a small peptide that forms a covalent, irreversible bond with its protein partner, SpyCatcher, allowing site-specific ligand coupling under physiological conditions. Inserting SpyTag into surface-exposed capsid sites enabled post-assembly functionalization of AAVs with SpyCatcher-fused targeting proteins. As proof-of-concept, we used SpyCatcher fusions with designed ankyrin repeat proteins (DARPins) specific for EGFR, EpCAM, and HER2. This conferred highly specific transduction of corresponding cancer cell lines with minimal off-target activity. Therapeutic potential was demonstrated by delivering a suicide gene, inducing selective cancer cell killing upon prodrug administration. This “one-fits-all” platform allows rapid and flexible retargeting without significantly altering the underlying vectors genome or production process. It supports the incorporation of large or complex ligands not amenable to genetic fusion and facilitates high-throughput preclinical evaluation strategies. By uniting capsid engineering with modular ligand display, our approach provides a scalable and versatile framework for precision gene delivery, broadening the applicability of rAAV in both therapeutic and discovery settings.

## Introduction

The 21st century has seen rapid advancements in gene therapeutics. Since the approval of Glybera, the first viral gene therapy in the US and Europe, at least 18 viral gene therapies have been marketed to treat cancer, rare monogenic diseases, hematological and neurological disorders, and more. These therapies leverage viruses’ natural capacity to deliver genetic material into target cells, enabling treatment at the genetic level *(1)*. Among viral vectors, adeno-associated viral (AAV) vectors have emerged as the leading platform for *in vivo* gene therapy, as demonstrated by seven approved gene therapies, including Zolgensma, Luxturna, and Beqvez (*2)*,(*3)*.

AAVs popularity stems from several key features, including its limited ability to integrate into the hosts genome (*4)*,(*5)*, their low immunogenicity and lack of pathogenicity (*6)* as well as the availability of serotypes with different tissue specificities (*7)*. Wild-type AAV (wtAAV) has a 4.7 kb single-stranded DNA genome flanked by inverted terminal repeats (ITRs) that function as packaging signals. The genome encodes genes for replication (*rep*) and structural capsid proteins (*cap*), including the proteins VP1, VP2, and VP3, which assemble into a 60-mer capsid (*8)*,(*9)*. While wtAAV infection can result in latent genome integration (*4)*,(*5)*, recombinant AAV (rAAV) vectors used in gene therapy remain episomal in transduced cells, minimizing the risk of insertional mutagenesis (*10)*.

Beyond natural AAV serotypes, engineered rAAV capsids are being developed for clinical applications by using genetic or chemical modifications and peptide insertions to achieve altered tissue specificities or capsid shielding (*11)*,(*12)*,(*13)*. AAV capsids can be functionalized by two primary strategies. Small peptides or protein domains are commonly inserted into exposed surface loops. In AAV2, position 587 in variable region VIII is the most utilized insertion site, enabling targeting of specific cell types by display of targeting ligands (*14)*,(*15)*,(*16)*. Insertions at this site also ablate natural tropism by disrupting primary receptor (heparan sulfate) binding (*17)*,(*18)*. Larger inserts, such as nanobodies for targeting cell-surface receptors, can be accommodated in variable region IV (*19)*. Alternatively, larger peptides or proteins can be fused to the N-terminus of VP2 and exposed through a pore in the viral capsid, allowing the display of affibodies, designed ankyrin repeat proteins (DARPins), or single-chain variable fragments (scFv) for precise targeting (*20)*,(*21)*.

While genetic fusion strategies have enabled progress in targeted AAV gene delivery, they face critical limitations. Each new target requires vector redesign, which is labor-intensive, results in a high variability of vectors and necessitates extensive characterization. Additionally, AAV capsids tolerate only limited insertions: surface-exposed loops and the VP2 N-terminus typically accommodate only peptides or small proteins, while larger proteins often destabilize the capsid or interfere with assembly. Furthermore, targeting ligands are restricted to proteins and molecules stable in the reducing environment of the host cell nucleus during vector production, excluding many antibody fragments and native ligands. Alternative strategies like bispecific antibodies (*22)* or chemical coupling strategies like biotinylation (*23)* often lack the covalent binding stability needed for *in vivo* applications or require additional chemical (*24)* or enzymatic steps (e.g. BirA-mediated biotinylation (*25)*).

The SpyTag/SpyCatcher system, derived from a split fragment of the *Streptococcus pyogenes* fibronectin-binding protein FbaB (*26)*, provides a compelling solution. This system enables site-specific covalent coupling of a peptide tag (SpyTag) with its protein partner (SpyCatcher) under physiological conditions without the need for cofactors or chemical catalysis. Previous studies have leveraged this system to equip viral particles post-assembly. For example, Kasaraneni *et al*. (*27)* inserted a SpyTag into the envelope of Sindbis-pseudotyped lentiviral vectors and redirected them to HER2+ cells using SpyCatcher fused to HER2-specific designed ankyrin repeat proteins (DARPins) or Fab fragments. Similarly, Kadkhodazadeh *et al*. (*28)* used SpyTag insertion into the HI loop of the adenovirus type-5 (Ad5) fiber knob to enable modular targeting via SpyCatcher-nanobody conjugates. More recently, Zhang *et al*. (*29)* equipped AAV2 with SpyTags by inserting an unnatural amino acid into VP3 and coupled the vector to SpyCatcher-fused nanobodies via a SpyTag-click-chemistry adapter, enabling targeted gene delivery *in vitro* and *in vivo*.

In this study, we present a genetically integrated SpyTag displayed directly in the AAV capsid, enabling covalent and modular redirection of rAAV2 via post-assembly coupling to SpyCatcher-ligand fusions. This approach unites the advantages of genetic engineering and modular post-production functionalization: a single, universal SpyTag-AAV vector that can be redirected to diverse cell types simply by exchanging the SpyCatcher-fused targeting ligand.

We demonstrate the flexibility of this platform by equipping SpyTag-AAVs with DARPins targeting EGFR, EpCAM and HER2, which successfully redirected the vector to corresponding cancer cell lines. Furthermore, we validated therapeutic functionality through the delivery of a suicide gene and subsequent prodrug-induced cell death. By decoupling targeting from vector production, our platform streamlines the rapid screening and preclinical evaluation of new AAV-based therapeutics. The ability to test diverse ligands without reengineering the capsid accelerates discovery and facilitates the development of receptor-specific gene delivery strategies.

## Results

### Design of a modular AAV-based transduction platform

Our transduction platform comprises an engineered recombinant AAV2 (rAAV2) and an exchangeable targeting ligand that mediates selective binding of a cellular receptor and thereby targeted transduction of cells (Fig. 1A). The AAV vector carries a mutation in its heparan sulfate proteoglycan (HSPG) binding motif, which disables primary receptor binding (*17)*,(*18)*, and displays a genetically inserted SpyTag peptide (*26)* on its capsid. The targeting ligand comprises a fusion protein of the SpyTag-binding partner SpyCatcher (*26)*,(*30)* and a designed ankyrin repeat protein (DARPin) specific for a cell surface receptor (*31)*. The SpyCatcher forms an amide bond with the SpyTag, thereby equipping the AAV with the DARPin. The DARPin mediates specific interaction with a cell surface receptor, resulting in selective transduction of the target cell.

**Figure 1:**
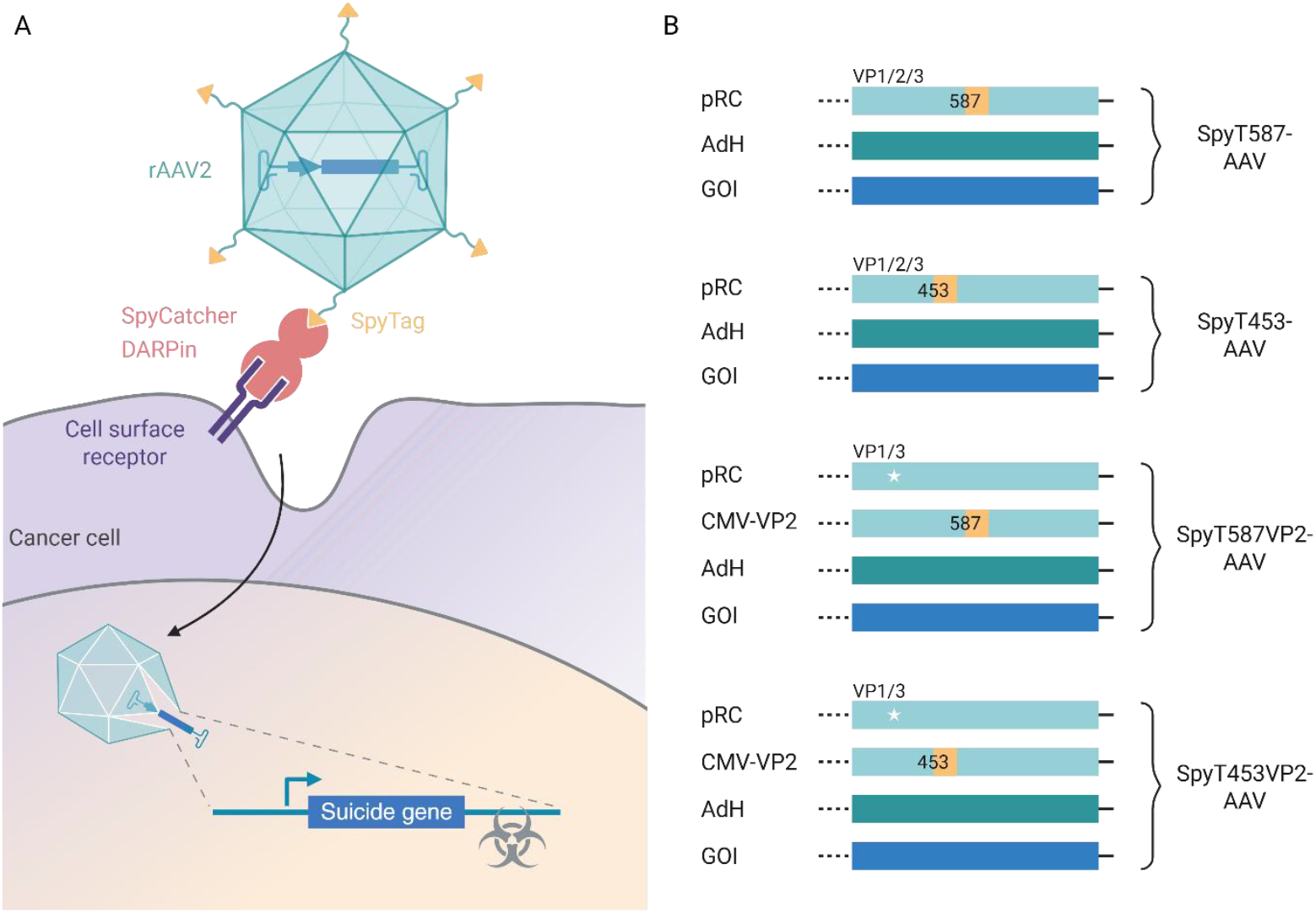
Design of a modular AAV-based transduction platform for cancer cell therapy. **A**: Schematic illustration of the modular AAV retargeting platform designed for cancer cell transduction. The rAAV2 capsid displays a genetically inserted SpyTag, which binds to a SpyCatcher-DARPin fusion protein. The DARPin binds to cancer cell surface receptors, enabling the delivery of a suicide gene that triggers cell death upon prodrug administration. **B**: Overview of SpyTag-AAV constructs used for AAV production. The SpyTag001 peptide (AHIVMVDAYKPTK) (yellow) was inserted at positions 587 or 453 of the viral capsid, either in all three viral capsid proteins (VP1, VP2, and VP3) or exclusively into VP2 with a pRC plasmid harboring a mutated VP2 start codon (white star) supplied *in trans*. Abbreviations: AdH, Adenovirus helper plasmid; DARPin, designed ankyrin repeat protein; GOI, gene of interest; pRC, rep-cap plasmid; rAAV, recombinant adeno-associated virus; SpyT, SpyTag.

### Implementation and characterization of the transduction platform

To implement and characterize the modular AAV-transduction platform, we selected DARPin E_01, a designed ankyrin repeat protein (DARPin) specific for the epidermal growth factor receptor (EGFR), which is commonly expressed on tumor cells (*32)*,(*33)*. DARPin E_01 has previously been utilized to retarget viral vectors, including AAVs, to EGFR-expressing cells (*34)*,(*35)*,(*36)*. We produced the SpyCatcher-DARPin_EGFR_ fusion protein (referred to hereafter as SpyC-DARPin_EGFR_) in *E. coli*, purified it using immobilized metal ion affinity chromatography (IMAC) and confirmed its integrity via SDS-PAGE and Coomassie staining (Fig. S1).

For the genetic integration of the SpyTag into the rAAV2 capsid, we used two sites: positions 587 in the variable region VIII (*14)* and 453 within the variable region IV (*37)*. Both positions are located in surface exposed loops of the AAV capsid and have previously demonstrated tolerance for peptide insertions for rational targeting of cells (*37)*,(*15)*,(*16)*, (*14)*,(*38)*. We inserted SpyTag at both positions in two configurations: (i) into all three viral capsid proteins (VP1, VP2 and VP3) using plasmids pMH321 and pHJW414 (see Table S4 and S5), and (ii) exclusively into VP2 using plasmids pHJW341 and pHJW351. For the latter, VP2 expression was driven by a CMV promoter *in trans* to a rep-cap plasmid harboring a mutated VP2 start codon (plasmid pHJW162), preventing production of unmodified VP2. To ablate the natural tropism of the AAV vectors and ensure transduction only in the presence of the DARPin, we introduced R585A and R588A mutations into the cap gene (*18)*,(*39)*. This process yielded four distinct AAV variants: SpyT587-AAV, SpyT453-AAV, SpyT587VP2-AAV, and SpyT453VP2-AAV (Fig. 1B).

SpyTag-AAVs carrying the fluorescent reporter gene mScarlet (plasmid CMV-mScarlet) were produced using a helper-free packaging system in HEK-293T cells and precipitated from cell culture supernatant using polyethylene glycol (PEG) (see Methods). Western blot analysis against the viral capsid proteins VP1, VP2 and VP3 confirmed the presence of the viral capsid proteins in all AAV preparations (Fig. 2).

**Figure 2:**
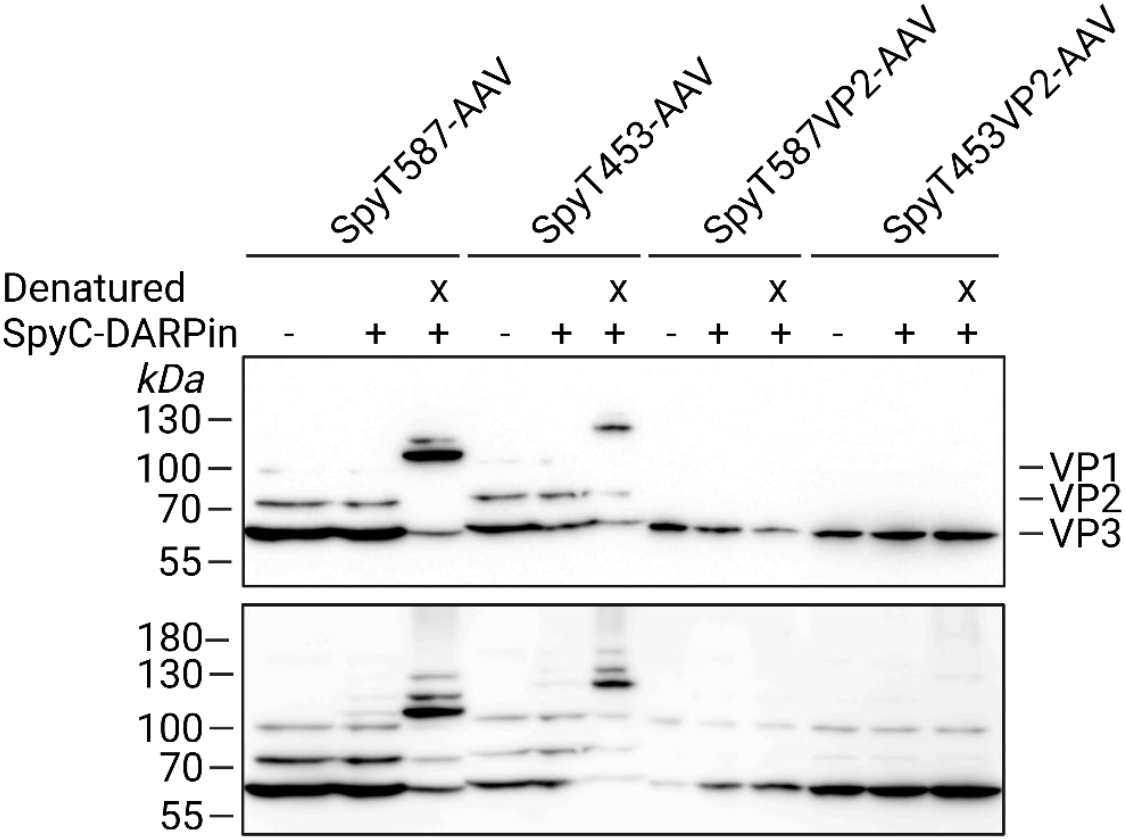
Evaluation of capsid composition and SpyCatcher-DARPin_EGFR_ coupling of SpyTag-AAVs. Western blot analysis of PEG-precipitated viral capsid proteins VP1, VP2 and VP3 of SpyT587-AAV, SpyT453-AAV, SpyT587VP2-AAV, and SpyT453VP2-AAV from cell culture supernatant. “+” indicates incubation with SpyCatcher-DARPin_EGFR_, “x” denotes AAV denaturation by boiling at 98 °C for 10 min before SC-DARPin coupling. The lower panel shows an overexposed version of the membrane presented in the upper panel, revealing weaker signals. SpyT-AAV and SpyC-DARPin_EGFR_ concentrations are listed in Table S1.

To assess SpyC-DARPin coupling, precipitated SpyT-AAVs were incubated with SpyC-DARPin either directly or following heat denaturation (98°C, 10 min), which served to expose any inaccessible SpyTags. Successful coupling was indicated by a molecular weight shift in capsid proteins on the Western blot due to the covalent attachment of the SpyC-DARPin.

Both SpyT587-AAV and SpyT453-AAV displayed all three viral proteins (Fig. 2 upper panel), while SpyT587VP2-AAV and SpyT453VP2-AAV capsids exhibited a weak VP2 signal, visible only after overexposure of the membrane (Fig. 2 lower panel). The latter two constructs rely on trans-complementation of SpyTag-modified VP2 which might disrupt the normal capsid stoichiometry of 1:1:10 for VP1:VP2:VP3 and could be the cause for poor incorporation of SpyTag-modified subunits. Similarly, SpyT587-AAV and SpyT453-AAV displayed additional higher molecular weight bands after SpyC-DARPin_EGFR_ incubation both in their native, albeit weak, and their denatured state while SpyT587VP2-AAV and SpyT453VP2-AAV only displayed weak additional bands after denaturation.

The low coupling efficiency in the native state with notable coupling to denatured capsid proteins confirms that SpyTags were correctly integrated but are likely buried within the intact capsid structure. This suggests that only a small subset of viral proteins expose functional SpyTags on the capsid surface.

To assess whether coupling SpyC-DARPin_EGFR_ to the SpyTag-AAVs enabled specific retargeting, we evaluated the vectors’ ability to transduce A-431 cells, which overexpress EGFR. To this aim, we assessed transduction efficiency at different SpyT-AAV doses and at different SpyC-DARPin protein concentrations. Serial dilutions of the vectors were pre-incubated with SpyC-DARPin_EGFR_ in cell culture medium before being added to the cells (Fig. 3). Flow cytometry analysis revealed that coupling of SpyC-DARPin_EGFR_ enabled targeting of SpyT-AAVs to target cells. The dilution series of SpyT-AAVs showed a vector-concentration dependent increase in mScarlet-positive cells, indicating successful transduction across all four capsid variants tested. Interestingly, despite the absence of detectable SpyC-DARPin_EGFR_ coupling in the Western blot analysis, SpyT587VP2-AAV and SpyT453VP2-AAV achieved transduction efficiencies comparable to SpyT587-AAV and SpyT453-AAV at the highest tested multiplicity of infection (MOIs). Notably, SpyT453-AAV maintained robust transduction even at MOIs as low as 190, underscoring its efficiency. Specificity was confirmed by the minimal off-target transduction observed in AAVs incubated without SpyC-DARPin_EGFR_ (< 6.2% for SpyT453-AAV and <1.2 % for all others at MOI 3.1*10^3^, Fig. 3), highlighting that the retargeting mechanism depends on the SpyC-DARPin_EGFR_ interaction.

**Figure 3:**
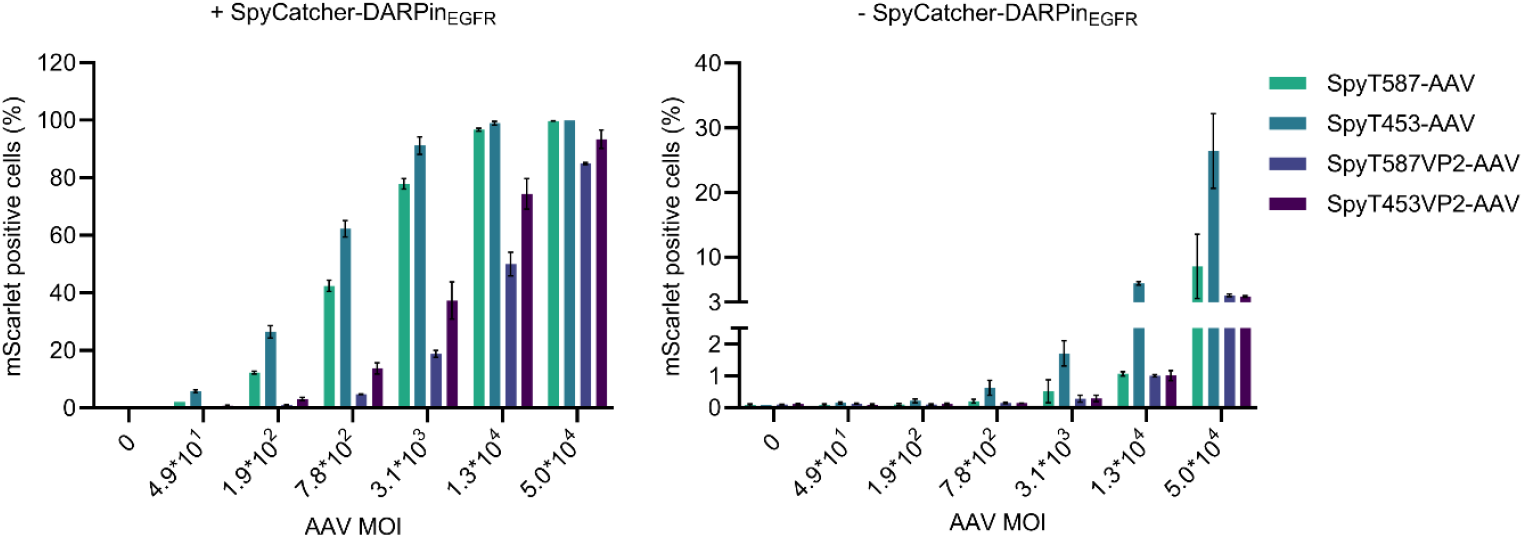
SpyCatcher-DARPin_EGFR_ specific targeting of EGFR-overexpressing A431 cells. Indicated SpyTag-AAV variants were evaluated for their transduction of EGFR-overexpressing A-431 cells with or without SpyCatcher-DARPin_EGFR_. AAVs were serially diluted (staring from 2.0*10^9^ vg/ml, then diluted serially 1:4) and each dilution was incubated with 100 nM of SpyC-DARPin_EGFR_ for 1 h at 37°C. Transduction mix was then added to A-431 cells and transduction efficiency was analyzed after 48-72 h by quantifying mScarlet positive cells by flow cytometry. Bars represent means ± SD of n=3 biological replicates.

Although integration site 587 has been widely utilized for peptide insertions into the AAV capsid, position 453 has not yet gained the same popularity (*40)*,(*9)*. Given that SpyT453-AAV demonstrated the highest transduction efficiency of all four AAVs, we selected SpyT453-AAV for subsequent experiments to further explore its potential in targeted gene delivery.

We next aimed to identify the optimal concentration and coupling conditions for SpyC-DARPin_EGFR_ to maximize SpyT453-AAV-mediated transduction. Serial dilutions of SpyC-DARPin_EGFR_ were incubated with SpyT453-AAV for 1 h at 37 °C or overnight at room temperature (Fig. 4A). Transduction of A-431 cells was efficient across a broad range of SpyC-DARPin_EGFR_ concentrations (1 µM to 1 nM). Peak transduction efficiency was observed at 10 nM SpyC-DARPin, with higher concentrations resulting in lower efficiencies, likely due to free SpyC-DARPin_EGFR_ blocking cell receptors. However, transduction efficiency declined sharply at concentrations below 1 nM, where SpyTag-SpyCatcher coupling seemed inefficient. Notably, temperature and incubation time had a minimal impact on transduction efficiency (Fig. 4A), therefore 1 h incubation at 37 °C was used for subsequent experiments.

**Figure 4:**
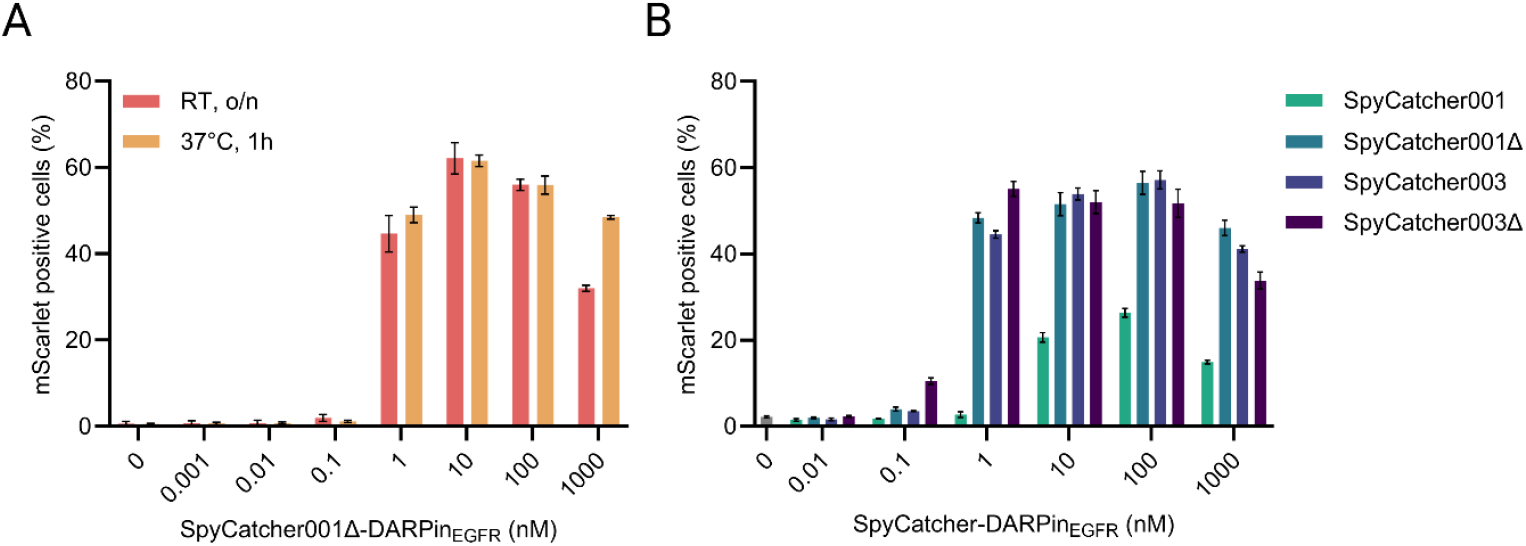
Influence of SpyCatcher coupling conditions and variants on SpyTag-AAV transduction efficiency. **A:** Titration of SpyCatcher-DARPin_EGFR_ concentration. SpyT453-AAV (3.1*10_7_ vg/ml, MOI: 7.8*10_2_) was pre-incubated with serial dilutions of SpyC-DARPin_EGFR_ for 1 h at 37 °C or overnight at room temperature before transduction of A-431 cells. **B:** Evaluation of SpyCatcher variants on SpyTag-AAV transduction efficiency. SpyT453-AAV (1.0*10^8^ vg/ml, MOI: 2.5*10_3_) was pre-incubated with serial dilutions of various SpyCatcher-DARPin_EGFR_ fusions for 1 h at 37 °C. Transduction efficiency was measured as mScarlet positive A-431 cells by flow cytometry. Bars represent means ± SD of *n*=3 biological replicates.

Thus far, we had employed the truncated SpyCatcher001ΔN3ΔC2 (*30)* (referred to hereafter as SpyCatcher001Δ) for coupling. SpyCatcher001Δ was chosen for its reported reduced interaction with cell surface receptors compared to SpyCatcher001 (*27)*. To explore whether alternative SpyCatcher variants could enhance transduction, we generated and tested DARPin_EGFR_ fusions with SpyCatcher001 (*26)*, SpyCatcher003, which harbors a stabilized loop and increased surface polarity for enhanced reaction compared to the first generation (*41)*, and our newly engineered variant, SpyCatcher003Δ, designed through sequence comparisons of SpyCatcher001 and SpyCatcher001Δ (Fig. S2 and S3). All variants enabled efficient A-431 cell transduction, but none considerably outperformed Spycatcher001Δ, except for SpyCatcher003Δ in one instance, showing a modest advantage at 1 nM (Fig. 4B). Since the SpyC-DARPin_EGFR_ construct did not critically influence overall transduction efficiency, we chose to continue using Spycatcher001Δ for subsequent experiments.

### Modular redirection of AAV to different cancer cells

To demonstrate that our AAV platform enables modular and versatile retargeting to different cell lines, we replaced the EGFR-specific DARPin E_01 with DARPin Ec1 (*42)* and DARPin 9_29 (*32)*, which specifically target EpCAM and Her2/ErbB2, respectively (Fig. S4).

SpyT453-AAV equipped with DARPin E01 selectively transduced A-431 and SK-OV-3 cells, both of which express EGFR (*43)*,(*44)* (Fig. 5). DARPin Ec1 enabled specific redirection to EpCAM-expressing MDA-MB-453 cells (*45)*,(*46)* and, to a lesser extent, to A-431 cells, which express low levels of EpCAM (*47)*. Finally, DARPin 9_29 efficiently redirected AAVs to HER2-expressing SK-OV-3 cells (*48)*,(*49)* as well as MDA-MB-453 cells, which are presumed to be HER2-positive (*50)*,(*51)*,(*52)*,(*53)*.

**Figure 5:**
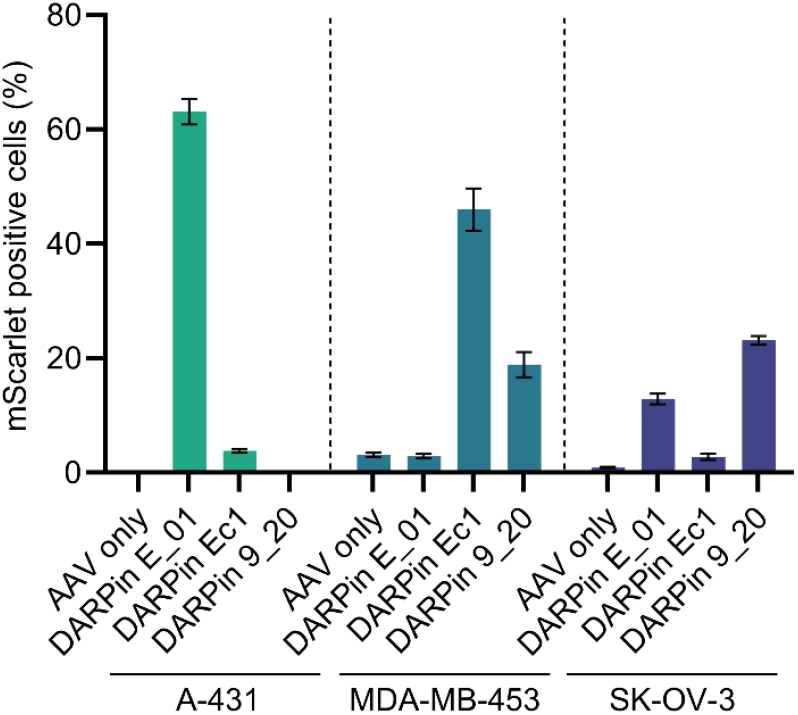
Modular retargeting of SpyTag-AAV to different cancer cell lines using surface marker-specific DARPins. SpyT453-AAV (1.4*10^8^ vg/ml, MOI: 3.5*10^3^) was pre-incubated with 10 nM of SpyC-DARPin fusions targeting EGFR (DARPin E_01), EpCAM (DARPin Ec1), Her2/ErbB2 (DARPin 9_20), or without targeting ligand (AAV only). A-431, MDA-MB-453 (EpCAM and HER2 expressing), and SK-OV-3 cells (Her2/ErbB2 expressing) were transduced. Transduction efficiency was determined by flow cytometry based on mScarlet expression. Bars represent means ± SD of *n*=3 biological replicates.

Overall, SpyT453-AAVs functionalized with SpyC-DARPin fusion proteins efficiently transduced target-expressing cancer cells while maintaining minimal off-target transduction in non-target expressing cell lines. These results underscore the flexibility of the platform in retargeting AAV vectors to diverse cell types via modular DARPin substitutions.

### Targeted killing of cancer cells using a suicide gene approach

As proof-of-concept for the modular retargeting, we applied our AAV system for viral-directed enzyme prodrug therapy (suicide gene therapy), wherein an exogenous enzyme is delivered into e.g. cancer cells, which then sensitizes the target cells to a non-toxic prodrug. Linamarase, an enzyme derived from cassava, has previously attracted attention for its anticancer properties and suicide gene potential (*54)*, (*55)*, (*56)*. Linamarase catalyzes conversion of the cyanoglucoside linamarin into cytotoxic hydrogen cyanide (HCN) (*57)*. Cyanide induces cell death by blocking oxidative phosphorylation in the mitochondrial respiratory chain (*58)*, (*59)* and kills surrounding cells through a bystander effect (*60)*, (*61)* often desired in the treatment of tumor sites.

To leverage our modular platform for viral directed enzyme prodrug therapy (VDEPT), we packaged the suicide gene linamarase into the SpyT453-AAV capsid. To this aim, the coding sequence for linamarase followed by a 2A peptide (T2A) was inserted upstream of the mScarlet reporter gene of plasmid CMV-mScarlet (resulting in plasmid pHJW427), allowing for co-expression of both genes. The resulting AAVs were characterized through Western blot analysis of the capsid proteins and by transduction efficiency assays in A-431 cells after coupling with SpyC-DARPin_EGFR_ (Fig. S5).

A-431 cells were transduced with SpyC-DARPin_EGFR_-coupled SpyT453-AAVs carrying either linamarase-mScarlet fusion or mScarlet alone as control before adding different concentrations of linamarin. Cell viability was evaluated using a Zombie violet live/dead staining by flow cytometry (Fig. 6). Additionally, hydrogen cyanide (HCN) production – a key indicator of linamarase activity – was quantified in the cell culture supernatant via gas chromatography mass spectrometry (GCMS) (Fig. 6). While the percentage of live cells of mScarlet control transduced cells remained relatively constant in the presence of increasing linamarin concentrations, a continuous decrease was evident for linamarase-mScarlet transduced cells. A significantly decreased live cell number was seen for linamarin concentrations between 1000 and 2000 µg/ml linamarin. Proportionally, the amount of HCN produced by cells increased with increasing linamarin concentrations in the presence of linamarase only, reaching up to 24 µg/ml HCN. This equals an average substrate conversion efficiency of approximately 10% (Table S3). Considering evaporation of HCN during the handling of the experiment, the actual conversion rate is likely to be higher. These experiments confirmed successful gene delivery and enzymatic activity, providing proof-of-concept for the use of SpyT453-AAVs in VDEPT applications.

**Figure 6:**
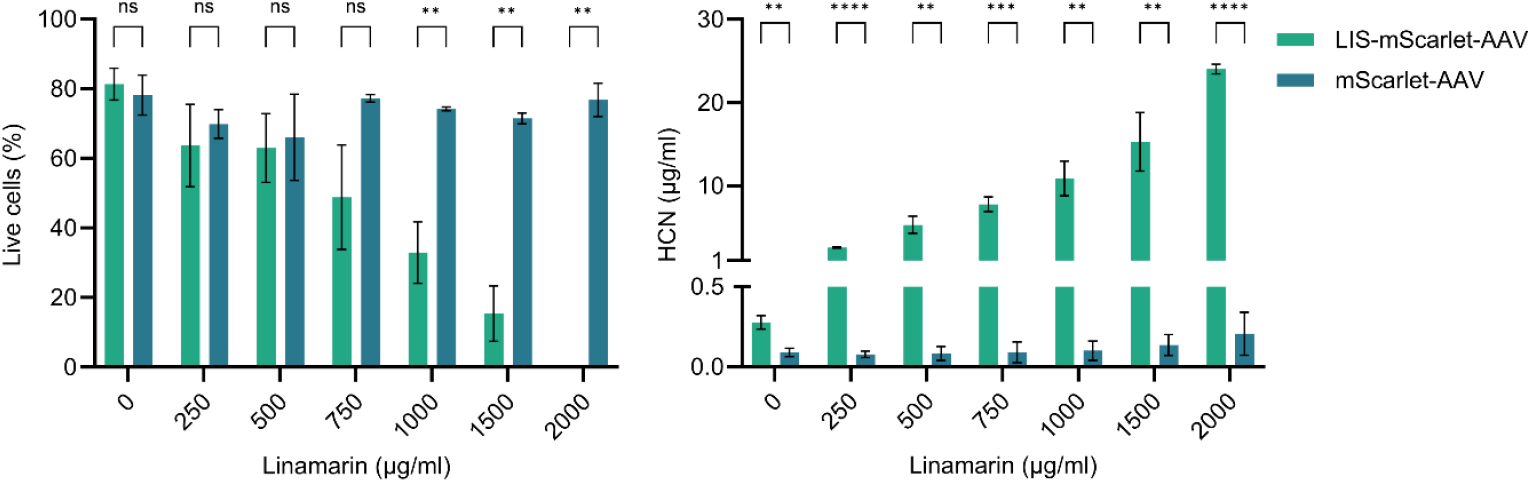
Targeted cancer cell killing using the linamarase suicide gene. Live cell quantification by flow cytometry with Zombie violet staining and HCN production analysis after delivering the linamarase suicide gene and incubation of cells with different concentrations of linamarin. SpyT453-AAV (2.4*10^8^ vg/ml, MOI: 5.9*10^3^) carrying either linamarase-mScarlet fusion or mScarlet gene alone (control) were pre-incubated with 10 nM SpyC-DARPin_EGFR_ and used to transduce A-431 cells. Statistical significance is indicated as follows: ns (*p* > 0.05), **(*p* ≤ 0.01), ***(*p* ≤ 0.001), ****(*p* ≤ 0.0001). Bars represent means ± SD of *n*=3 biological replicates.

## Discussion

Recombinant adeno-associated viral vectors have emerged as the vector of choice for *in vivo* gene delivery, but their widespread adoption faces key challenges. These include limited tissue specificity, manufacturing challenges, and the presence of neutralizing antibodies in patients (*62)*. Numerous studies have proposed innovative solutions to address these hurdles, with modular targeting platforms showing promise for improved specificity and simplified production for discovery and preclinical studies.

In this study, we developed a modular AAV-based retargeting platform, for the first time leveraging the SpyTag/SpyCatcher technology via genetic integration for the rapid and flexible post-assembly functionalization of AAV capsids with targeting ligands. By inserting SpyTags into defined positions on the AAV2 capsid, we created a universal vector scaffold that can be redirected to diverse targets by coupling to SpyCatcher-fusion ligands. We demonstrated this modularity by equipping the vector with different SpyCatcher-DARPin fusion proteins specific for EGFR, EpCAM, and HER2/ErbB2, thereby achieving efficient and specific transduction of corresponding cancer cell lines. Additionally, we validated the platform’s therapeutic functionality by delivering a suicide gene, which induced selective cell death upon prodrug activation.

Our approach builds on earlier strategies for modular AAV targeting, including the split-intein-based approach (*21)*, where the surface of fully assembled AAV capsids carrying a C-intein at the N-terminus of VP2 were equipped with N-intein-fused single-chain variable fragments (scFvs) or DARPins and the click-chemistry-based approach for SpyTag-display on AAV (*29)*. Unlike the latter method, our system relies solely on genetic incorporation of the SpyTag into the capsid, providing a simpler and more direct route to post-assembly vector modification. We identified capsid positions 453 and 587 as suitable for SpyTag insertion and showed that incorporation into all three capsid proteins led to more efficient surface display than selective VP2-only modification - likely due to better incorporation ratios and capsid assembly dynamics.

Although Western blot analysis suggested that only a subset of capsid proteins displayed accessible SpyTags in the native conformation, even limited functionalization was sufficient to achieve high transduction efficiencies. This suggests that only a few accessible ligands may be needed for effective receptor-mediated entry. Functionalization could potentially be increased by the introduction of linker sequences or modifying flanking residues to enhance externalization, as suggested in previous studies (*63)*. Simultaneously, this low functionalization enables the application of the system with a minimum of non-human-derived protein adapters, thereby lowering the risk of immunogenic reaction. Furthermore, the use of minimally modified rAAV2 capsids ensures compatibility with standard production workflows, while the SpyTag/SpyCatcher system’s robustness across physiological buffers simplifies application in biologically relevant settings. Compared to intein-based or chemically modified systems, our approach avoids the need for reducing agents or non-physiological conditions and expands the range of compatible ligands, including disulfide-rich or structurally complex proteins.

The successful coupling of a newly engineered SpyCatcher variant (SpyCatcher003Δ) further demonstrates the flexibility of the system and points to opportunities for optimizing binding kinetics and efficiency through next-generation SpyTag/SpyCatcher pairs. The modular post-assembly nature of this platform also enables future exploration of complex ligands such as scFvs, nanobodies, or native ligands (*21)*,(*27)* or multi-specific targeting strategies, for example by coupling bispecific DARPins (*64)* or ligand mixtures to enhance cell-type specificity and therapeutic precision.

Beyond its therapeutic potential, this platform offers a valuable tool for the rapid screening and preclinical evaluation of targeted AAVs. The ability to use a single capsid scaffold and test multiple targeting ligands in parallel accelerates the discovery pipeline and reduces the need for repeated vector engineering and validation. Furthermore, the use of a single capsid for various targets could justify the generation of a producer cell line and thereby save costs and facilitate the production process. While the bacterial origin of the SpyCatcher protein raises questions about immunogenicity, previous studies have reported limited immune responses in therapeutic contexts *(27), (65)*, though further evaluation will be essential in future translational work.

In conclusion, we present a genetically encoded, modular AAV platform that enables post-assembly functionalization via the SpyTag/SpyCatcher system. This strategy allows for flexible, efficient redirection of AAVs to various cellular targets using interchangeable ligands, supporting both therapeutic applications and preclinical vector discovery with minimal need for capsid redesign.

## Materials and Methods

### Cloning of plasmids

Plasmids used and generated are listed in Table S4. Gene sequences were amplified using polymerase chain reaction (PCR) and assembled by Gibson cloning (*66)* or restriction enzyme cloning. Mutations and SpyTag sequences were introduced using oligonucleotides and polymerase chain reaction (PCR).

### Protein production and purification

For the production of all SpyCatcher-DARPin proteins, the respective plasmids were transformed into *E. coli* BL21 Star (DE3) (Thermo Fisher Scientific, Cat. No. C601003). Transformed bacteria were selected in LB medium supplemented with ampicillin (100 µg/ml) and grown at 37 °C and 150 rpm to an OD_600_ of 0.7, at which expression was induced with 1 mM isopropyl-*β*-D-thiogalactopyranoside (IPTG). After incubation at 30 °C and 150 rpm for 4 h, bacteria were harvested by centrifugation at 6,000 x *g* for 10 min. The cell pellet was resuspended in lysis buffer [50 mM NaH_2_PO_4_, 300 mM NaCl, 10 mM imidazole, pH 8.0], shock frozen in liquid nitrogen, and stored at -80 °C. For purification, frozen pellets were thawed at 37 °C in a water bath and lysed by sonication (Bandelin Sonopuls HD3100, 10 min with 60% amplitude, 0.5 s pulse and 1 s pause intervals). Following clarification of the lysate by centrifugation at 30,000 x g for 30 min at 4 °C, the supernatant was loaded onto a gravity flow column with Ni-NTA Superflow agarose (Qiagen, Cat. No. 30430). The column was washed with 20 column volumes (CV) of wash buffer [50 mM NaH_2_PO_4_, 300 mM NaCl, 20 mM imidazole, pH 8.0], and the purified protein was eluted in 6 CV elution buffer [50 mM NaH_2_PO_4_, 300 mM NaCl, 250 mM imidazole, pH 8.0]. Afterwards, the purified protein was dialyzed against PBS [2.7 mM KCl, 1.5 mM KH_2_PO_4_, 8.1 mM Na_2_HPO_4_, and 137 mM NaCl (pH 7.4)] using 10 K MWCO dialysis tubing (Thermo Fisher Scientific, Cat. No. 88243). Protein aliquots were shock frozen in liquid nitrogen and stored at -80 °C.

### Protein characterization

Protein identity and purity was analyzed by sodium dodecyl sulfate-polyacrylamide gel electrophoresis (SDS-PAGE) using 10% gels followed by Coomassie staining. As protein size marker, PageRuler prestained protein ladder (Thermo Fisher Scientific, Cat. No. 26616) was used. Protein concentrations were determined by Bradford assay (Bio-Rad, Cat. No. 500-0006) using bovine serum albumin (Sigma-Aldrich, Cat. No. 05479) as standard.

### Cell culture

HEK-293T (German Collection of Microorganisms and Cell Cultures (DSMZ), Cat. No. ACC 635) and A-431 (DSMZ, Cat. No. ACC 91) cells were cultivated in Dulbecco’s modified Eagle’s medium (DMEM) (PAN Biotech, Cat. No. P04-03550), supplemented with 10% (v/v) fetal bovine serum (FBS) (PAN Biotech, Cat. No. P30-3602, Lot. No. P150702), 100 U/ml penicillin, and 100 μg/ml streptomycin. SK-OV-3 cells (ATCC, Cat. No. HTB-77) were maintained in McCoy’s 5A medium (Sigma-Aldrich, catalog no. M8403) supplemented with 10% (v/v) FBS, 2 mM L-glutamine (Thermo Fisher Scientific, Cat. No. 25030-024), 100 U/ml penicillin, and 100 μg/ml streptomycin. MDA-MB-453 cells (ATCC, Cat. No. HTB-131) were cultivated in RPMI 1640 medium (Thermo Fisher Scientific, Cat. No. 61870-010) supplemented with 10% (v/v) FBS, 100 U/ml penicillin, and 100 μg/ml streptomycin. All cells were cultivated at 37 °C in a humidified atmosphere containing 5% CO_2_ and passaged upon reaching approximately 80% confluence.

### AAV production and purification

AAV vectors were produced using the adenovirus helper-free packaging system (*67)*. For the production of SpyTag-AAVs, HEK-293T cells were transfected with an AAV2 rep-cap plasmid (Cell Biolabs (Cat.No. VPK-402)), an adenovirus helper plasmid (Cell Biolabs (Cat.No. VPK-402)) and a self-complementary vector plasmid pCMV-mScarlet (or derivatives of it) (a gift from Dirk Grimm) in an equimolar ratio. For AAV production, 8*10^6^ HEK-293T cells were seeded per 15-cm cell culture dish and 24 h later, transfected with 60 µg of total plasmid DNA combined with 200 µg of polyethylenimine (PEI, MW 25,000) in 3 ml OptiMEM (Thermo Fisher Scientific, Cat. No. 22600-134). After 72 h, AAVs were precipitated from the supernatant using polyethylenglycole (PEG). To this end, the supernatant was collected and combined with a 40% PEG 8000 solution [40% (w/v) PEG 8000, 0.41 M NaCl, pH 7.4] to a final PEG concentration of 8% and incubated at 4 °C with slight agitation overnight. AAVs were then harvested by centrifugation at 4 °C and 2,828 x *g* for 15 min. The pellet was resuspended in an appropriate amount of sterile PBS (Thermo Fisher Scientific, Cat. No. 14190094) and filtered through a 0.45 µm polyvinylidene difluoride (PVDF) syringe filter. Aliquots were shock-frozen in liquid nitrogen and stored at -80 °C.

### AAV characterization and quantification

AAV preparations were analyzed for the presence of capsid proteins and SpyCatcher coupling by Western blotting against the viral capsid proteins. Depending on the condition, AAVs were either combined with SpyCatcher-DARPin protein, denatured at 98 °C for 10 min and then combined with SpyCatcher-DARPin protein, or combined with PBS and incubated at room temperature for 1 h. Samples were then mixed with 5x SDS loading buffer [50% (v/v) glycerol, 0.3125 M Tris-HCL pH 6.8, 0.05 % (w/v) bromophenol blue, 10% (w/v) SDS, 12.5% (v/v) 2-mercaptoethanol] and incubated at 70 °C for 10 min. AAV and protein concentrations are listed in Table S1. After separating the samples by SDS-PAGE, the proteins were blotted onto a methanol-activated PVDF membrane. The membrane was then blocked in blocking buffer [PBS containing 5% (w/v) low fat milk powder (Carl Roth, blotting grade, Cat. No. T145.3)] for 1 h at room temperature before incubation with anti-AAV VP1/VP2/VP3 B1 antibody (Progen, Cat. No. 65158) diluted 1:100 in binding buffer [PBS containing 2.5% (w/v) milk powder] at 4 °C overnight. Subsequently, the membrane was washed with PBS-T [PBS supplemented with 0.05% (v/v) Tween-20] and incubated with a secondary anti-mouse horseradish peroxidase (HRP)-conjugated antibody (Amersham Cytiva, Cat. No. NA931) diluted 1:5000 in binding buffer. After washing with PBS-T, either Pierce ECL Western Blotting-Substrate (Thermo Fisher Scientific, Cat. No. 32109) (Figure 2, upper panel) or SuperSignal West Femto Maximum Sensitivity Substrate (Thermo Fisher Scientific, Cat. No. 34094) (Figure 2, lower panel, Fig. S5) was added and chemiluminescence was imaged using the ImageQuant LAS 4000 Mini System (GE Healthcare).

The genomic titer of AAV vectors was determined by quantitative real-time PCR (qPCR). AAV samples were DNase I (New England Biolabs, Cat. No. M0303S) digested at 37 °C for 30 min. Then, serial dilutions of AAVs and of the vector plasmid pCMV-mScarlet as standard were prepared, and oligonucleotides 5’-TGCCCAGTACATGACCTTATGG-3’ and 5’-GAAATCCCCGTGAGTCAAACC-3’ were used to amplify a 134 bp fragment from the CMV promoter using the PowerTrack SYBR Green Mastermix (Thermo Fisher Scientific, Cat. No. A46109). The qPCR was performed on a CFX384 thermocycler (Bio-Rad) with the following temperature protocol: 3 min at 95 °C followed by 40 cycles of 15 s at 95 °C, 30 s at 60 °C, read plate, finally followed by a melting curve.

### AAV transduction experiments

For transduction experiments, 4,000 cells were seeded per well of a 96-well plate. After 24 h, SpyTag-AAVs were mixed with SpyCatcher-DARPin protein in transduction medium (DMEM, supplemented with 10% (v/v) FBS, 100 U/ml penicillin,100 μg/ml streptomycin and 10 mM HEPES (Thermo Fisher Scientific, Cat. No. 15630056)) and incubated at 37 °C for 1 h or as indicated. The final transduction mix consisted of 10 µl appropriately diluted AAV and 10 µl appropriately diluted SpyCatcher-DARPin protein, filled up to a final volume of 100 µl with transduction medium. Following complete removal of medium from the cells, 100 µl of transduction mix was added per well and cells were incubated at 37 °C in a humidified atmosphere containing 5% CO_2_ for 48-72 h.

For linamarin experiments, serial dilutions of linamarin (α-hydroxyisobutyronitrile β-D-glucopyranoside, Merck, Cat. No. 68264) in ddH_2_O were prepared. 20 µl linamarin dilutions were added to cells 24 h after transduction (final volume: 120 µl per well). The 96-well plate was then sealed with qPCR covering film (Greiner, Cat. No. 676040) in order to prevent gaseous HCN from evaporating or spreading to other wells and incubated at 37 °C in a humidified atmosphere containing 5% CO_2_ for 48 h.

### Flow cytometry

For analysis by flow cytometry, cells were washed with PBS and detached by the addition of 50 µl of trypsin/EDTA solution (PAN Biotech, Cat. No. P10-023100) per well. Then, 200 µl of FACS buffer (PBS supplemented with 2% (v/v) FBS) were added per well, cells were resuspended and analyzed by flow cytometry. For linamarin experiments with live/dead analysis, cells were stained using the Zombie Violet Fixable Viability Kit (Biolegend, Cat. No. 423113) according to the standard cell staining protocol. In brief, following resuspension in FACS buffer, cells were washed with PBS and resuspended in 100 µl staining solution containing 1:500 Zombie Violet dye in PBS. After incubation for 30 min at RT in the dark, cells were washed with FACS buffer and finally resuspended in 220 µl of FACS buffer. Cells were analyzed for mScarlet transgene expression and Zombie Violet staining using an Attune NxT flow cytometer (Thermo Fisher Scientific). BFP and Zombie Violet were excited with a 405-nm laser and detected using a 440/50-nm filter, while mScarlet was excited using a 561-nm laser and detected using a 620/15-nm filter. Cellular autofluorescence was measured in the unused BFP channel for experiments without Zombie Violet or using a 637-nm laser and 670/14-nm emission filter for experiments with Zombie Violet staining. Flow cytometry data was analyzed using FlowJo (v10.9.0, Becton Dickinson) with the gating strategy depicted in Figures S6 and S7.

### Head space-Solid phase microextraction Gas chromatography mass spectrometry (HS-SPME GCMS)

The quantification of HCN in samples was performed with a single quadrupole GC-MS system QP2010 SE (Shimadzu, Japan) equipped with a PAL autosampler AOC 5000 (CTC Analytics, Zwingen, Switzerland). The HS SPME technique was used for sample preparation. For this purpose, 200 ± 1 mg Na_2_SO4, 400 µl sample or standard solution, 20 µl of a 3 µg/ml CD3CN solution as internal standard and 75 µl of 85 % phosphoric acid were added to each HS vial and immediately sealed with PTFE/silicone septa. HS-SPME was achieved on a CBX/PDMS fiber (75 µm thickness) with 10 minutes extraction time at 35 °C with agitation followed by thermal desorption directly into the splitless injector at 250 °C. After each run, the fiber was conditioned at 250 °C for 30 minutes. A He column flow of 1 ml/min was realized, which corresponds to a linear velocity of 36 cm/s. The following temperature program was carried out on a DB-FFAP column (30 m×0.25 mm, thickness 0.25 µm): start temperature 40 °C for 8 min, heating with 40 K/min to 240 °C and 5 min hold time at final temperature of 240 °C. The mass spectra were recorded in SIM mode at m/z of 27 and 44. The MS interface was set to 250 °C and the ion source temperature to 200 °C. Internal standard calibration was done between 0 and 25 µg/ml HCN, achieving correlation coefficients of 99.95 %.

### Statistical analysis

For the analysis of differences between Linamarase-mScarlet-carrying AAV and mScarlet control AAV, unpaired two-sided t tests under no assumption of consistent standard deviations with correction for multiple comparisons (Holm-Sidak method) were performed using GraphPad Prism.

### Software

Flow cytometry data was analyzed with FlowJo (v10.9.0, Becton, Dickinson and Company). Data was analyzed and plotted using GraphPad Prism (v10.2.0, GraphPad Software). Figure 1 was created with BioRender.

## Acknowledgements

We acknowledge the excellent scientific and technical assistance of the Signalling Factory Core Facility staff of the University of Freiburg for help on flow cytometry and providing cell lines. We are grateful to Dirk Grimm (Heidelberg University, Germany) for providing the plasmid pCMV-mScarlet. This work was supported by the Deutsche Forschungsgemeinschaft (DFG, German Research Foundation) under Germany’s Excellence Strategy CIBSS, EXC-2189, Project ID: 390939984.

The authors declare no competing financial interest.

During the preparation of this work the authors used ChatGPT in order to improve readability and language. After using this tool, the authors reviewed and edited the content as needed and take full responsibility for the content of the publication.

## References

(1.) Chancellor, D., Barrett, D., Nguyen-Jatkoe, L., Millington, S., and Eckhardt, F. (2023) The state of cell and gene therapy in 2023, Molecular Therapy 31, 3376–3388.

(2.) Wang, J.-H., Gessler, D. J., Zhan, W., Gallagher, T. L., and Gao, G. (2024) Adeno-associated virus as a delivery vector for gene therapy of human diseases, Signal Transduction and Targeted Therapy 9, 78.

(3.) Dhillon, S. (2024) Fidanacogene Elaparvovec: First Approval, Drugs 84, 479–486.

(4.) Kotin, R. M., Siniscalco, M., Samulski, R. J., Zhu, X. D., Hunter, L., Laughlin, C. A., McLaughlin, S., Muzyczka, N., Rocchi, M., and Berns, K. I. (1990) Site-specific integration by adeno-associated virus, Proceedings of the National Academy of Sciences 87, 2211–2215.

(5.) Meneses, P., Berns Kenneth, I., and Winocour, E. (2000) DNA Sequence Motifs Which Direct Adeno-Associated Virus Site-Specific Integration in a Model System, Journal of Virology 74, 6213–6216.

(6.) Zaiss, A. K., and Muruve, D. A. (2008) Immunity to adeno-associated virus vectors in animals and humans: a continued challenge, Gene Therapy 15, 808–816.

(7.) Issa, S. S., Shaimardanova, A. A., Solovyeva, V. V., and Rizvanov, A. A. (2023) Various AAV Serotypes and Their Applications in Gene Therapy: An Overview, In Cells.

(8.) Weitzman, M. D., and Linden, R. M. (2011) Adeno-Associated Virus Biology, In Adeno-Associated Virus: Methods and Protocols (Snyder, R. O., and Moullier, P., Eds.), pp 1–23, Humana Press, Totowa, NJ.

(9.) Wagner, H. J., Weber, W., and Fussenegger, M. (2021) Synthetic Biology: Emerging Concepts to Design and Advance Adeno-Associated Viral Vectors for Gene Therapy, Advanced Science 8, 2004018.

(10.) Smith, R. H. (2008) Adeno-associated virus integration: virus versus vector, Gene Therapy 15, 817–822.

(11.) Kanaan, N. M., Sellnow, R. C., Boye, S. L., Coberly, B., Bennett, A., Agbandje-McKenna, M., Sortwell, C. E., Hauswirth, W. W., Boye, S. E., and Manfredsson, F.P. (2017) Rationally Engineered AAV Capsids Improve Transduction and Volumetric Spread in the CNS, Molecular Therapy -Nucleic Acids 8, 184–197.

(12.) Wang, D., Li, S., Gessler, D. J., Xie, J., Zhong, L., Li, J., Tran, K., Van Vliet, K., Ren, L., Su, Q., He, R., Goetzmann, J. E., Flotte, T. R., Agbandje-McKenna, M., and Gao, G. (2018) A Rationally Engineered Capsid Variant of AAV9 for Systemic CNS-Directed and Peripheral Tissue-Detargeted Gene Delivery in Neonates, Molecular Therapy -Methods & Clinical Development 9, 234–246.

(13.) Lee, G. K., Maheshri, N., Kaspar, B., and Schaffer, D. V. (2005) PEG conjugation moderately protects adeno-associated viral vectors against antibody neutralization, Biotechnology and Bioengineering 92, 24–34.

(14.) Girod, A., Ried, M., Wobus, C., Lahm, H., Leike, K., Kleinschmidt, J., Deléage, G., and Hallek, M. (1999) Genetic capsid modifications allow efficient re-targeting of adeno-associated virus type 2, Nature Medicine 5, 1052–1056.

(15.) Nicklin, S. A., Buening, H., Dishart, K. L., de Alwis, M., Girod, A., Hacker, U., Thrasher, A. J., Ali, R. R., Hallek, M., and Baker, A. H. (2001) Efficient and Selective AAV2-Mediated Gene Transfer Directed to Human Vascular Endothelial Cells, Molecular Therapy 4, 174–181.

(16.) Ried Martin, U., Girod, A., Leike, K., Büning, H., and Hallek, M. (2002) Adeno-Associated Virus Capsids Displaying Immunoglobulin-Binding Domains Permit Antibody-Mediated Vector Retargeting to Specific Cell Surface Receptors, Journal of Virology 76, 4559–4566.

(17.) Kern, A., Schmidt, K., Leder, C., Müller, O. J., Wobus, C. E., Bettinger, K., Von der Lieth, C. W., King, J. A., and Kleinschmidt, J. A. (2003) Identification of a Heparin-Binding Motif on Adeno-Associated Virus Type 2 Capsids, Journal of Virology 77, 11072–11081.

(18.) Opie Shaun, R., Warrington, K. H., Agbandje-McKenna, M., Zolotukhin, S., and Muzyczka, N. (2003) Identification of Amino Acid Residues in the Capsid Proteins of Adeno-Associated Virus Type 2 That Contribute to Heparan Sulfate Proteoglycan Binding, Journal of Virology 77, 6995–7006.

(19.) Eichhoff, A. M., Börner, K., Albrecht, B., Schäfer, W., Baum, N., Haag, F., Körbelin, J., Trepel, M., Braren, I., Grimm, D., Adriouch, S., and Koch-Nolte, F. (2019) Nanobody-Enhanced Targeting of AAV Gene Therapy Vectors, Molecular Therapy - Methods & Clinical Development 15, 211–220.

(20.) Nord, K., Gunneriusson, E., Ringdahl, J., Ståhl, S., Uhlén, M., and Nygren, P.-Å. (1997) Binding proteins selected from combinatorial libraries of an α-helical bacterial receptor domain, Nature Biotechnology 15, 772–777.

(21.) Muik, A., Reul, J., Friedel, T., Muth, A., Hartmann, K. P., Schneider, I. C., Münch, R. C., and Buchholz, C. J. (2017) Covalent coupling of high-affinity ligands to the surface of viral vector particles by protein trans-splicing mediates cell type-specific gene transfer, Biomaterials 144, 84–94.

(22.) Bartlett, J. S., Kleinschmidt, J., Boucher, R. C., and Samulski, R. J. (1999) Targeted adeno-associated virus vector transduction of nonpermissive cells mediated by a bispecific F(ab’γ)2 antibody, Nature Biotechnology 17, 181–186.

(23.) Stachler, M. D., Chen, I., Ting, A. Y., and Bartlett, J. S. (2008) Site-specific Modification of AAV Vector Particles With Biophysical Probes and Targeting Ligands Using Biotin Ligase, Molecular Therapy 16, 1467–1473.

(24.) Lee, S., and Ahn, H. J. (2019) Anti-EpCAM-conjugated adeno-associated virus serotype 2 for systemic delivery of EGFR shRNA: Its retargeting and antitumor effects on OVCAR3 ovarian cancer in vivo, Acta Biomaterialia 91, 258–269.

(25.) Barry, M. A. K. C. S.,, Debadyuti, G. E. A. K.,, Hoyin, M. T. M. G.,, and and Parrott, M. B. (2003) Biotinylated gene therapy vectors, Expert Opinion on Biological Therapy 3, 925–940.

(26.) Zakeri, B., Fierer, J. O., Celik, E., Chittock, E. C., Schwarz-Linek, U., Moy, V. T., and Howarth, M. (2012) Peptide tag forming a rapid covalent bond to a protein, through engineering a bacterial adhesin, Proceedings of the National Academy of Sciences 109, E690–E697.

(27.) Kasaraneni, N., Chamoun-Emanuelli Ana, M., Wright, G., and Chen, Z. (2017) Retargeting Lentiviruses via SpyCatcher-SpyTag Chemistry for Gene Delivery into Specific Cell Types, mBio 8, 10.1128/mbio.01860-01817.

(28.) Kadkhodazadeh, M., Mohajel, N., Behdani, M., Baesi, K., Khodaei, B., Azadmanesh, K., and Arashkia, A. (2022) Fiber manipulation and post-assembly nanobody conjugation for adenoviral vector retargeting through SpyTag-SpyCatcher protein ligation, Frontiers in Molecular Biosciences Volume 9 - 2022.

(29.) Zhang, Y., Chen, Z., Wang, X., Yan, R., Bao, H., Chu, X., Guo, L., Wang, X., Li, Y., Mu, Y., He, Q., Zhang, L., Zhang, C., Zhou, D., and Ji, D. (2024) Site-specific tethering nanobodies on recombinant adeno-associated virus vectors for retargeted gene therapy, Acta Biomaterialia 187, 304–315.

(30.) Li, L., Fierer, J. O., Rapoport, T. A., and Howarth, M. (2014) Structural Analysis and Optimization of the Covalent Association between SpyCatcher and a Peptide Tag, Journal of Molecular Biology 426, 309–317.

(31.) Plückthun, A. (2015) Designed Ankyrin Repeat Proteins (DARPins): Binding Proteins for Research, Diagnostics, and Therapy, Annual Review of Pharmacology and Toxicology 55, 489–511.

(32.) Steiner, D., Forrer, P., and Plückthun, A. (2008) Efficient Selection of DARPins with Sub-nanomolar Affinities using SRP Phage Display, Journal of Molecular Biology 382, 1211–1227.

(33.) Boersma, Y. L., Chao, G., Steiner, D., Wittrup, K. D., and Plückthun, A. (2011) Bispecific Designed Ankyrin Repeat Proteins (DARPins) Targeting Epidermal Growth Factor Receptor Inhibit A431 Cell Proliferation and Receptor Recycling*, Journal of Biological Chemistry 286, 41273–41285.

(34.) Dreier, B., Honegger, A., Hess, C., Nagy-Davidescu, G., Mittl, P. R. E., Grütter, M. G., Belousova, N., Mikheeva, G., Krasnykh, V., and Plückthun, A. (2013) Development of a generic adenovirus delivery system based on structure-guided design of bispecific trimeric DARPin adapters, Proceedings of the National Academy of Sciences 110, E869–E877.

(35.) Hanauer, J. R. H., Koch, V., Lauer, U. M., and Mühlebach, M. D. (2019) High-Affinity DARPin Allows Targeting of MeV to Glioblastoma Multiforme in Combination with Protease Targeting without Loss of Potency, Molecular Therapy - Oncolytics 15, 186–200.

(36.) Hörner, M., Jerez-Longres, C., Hudek, A., Hook, S., Yousefi, O. S., Schamel, W. W. A., Hörner, C., Zurbriggen, M. D., Ye, H., Wagner, H. J., and Weber, W. (2021) Spatiotemporally confined red light-controlled gene delivery at single-cell resolution using adeno-associated viral vectors, Science Advances 7, eabf0797.

(37.) Boucas, J., Lux, K., Huber, A., Schievenbusch, S., von Freyend, M. J., Perabo, L., Quadt-Humme, S., Odenthal, M., Hallek, M., and Büning, H. (2009) Engineering adeno-associated virus serotype 2-based targeting vectors using a new insertion site-position 453-and single point mutations, The Journal of Gene Medicine 11, 1103–1113.

(38.) Thadani, N. N., Yang, J., Moyo, B., Lee, C. M., Chen, M. Y., Bao, G., and Suh, J. (2020) Site-Specific Post-translational Surface Modification of Adeno-Associated Virus Vectors Using Leucine Zippers, ACS Synthetic Biology 9, 461–467.

(39.) Wu, P., Xiao, W., Conlon, T., Hughes, J., Agbandje-McKenna, M., Ferkol, T., Flotte, T., and Muzyczka, N. (2000) Mutational Analysis of the Adeno-Associated Virus Type 2 (AAV2) Capsid Gene and Construction of AAV2 Vectors with Altered Tropism, Journal of Virology 74, 8635–8647.

(40.) Büning, H., and Srivastava, A. (2019) Capsid Modifications for Targeting and Improving the Efficacy of AAV Vectors, Molecular Therapy Methods & Clinical Development 12, 248–265.

(41.) Keeble, A. H., Turkki, P., Stokes, S., Khairil Anuar, I. N. A., Rahikainen, R., Hytönen, V. P., and Howarth, M. (2019) Approaching infinite affinity through engineering of peptide–protein interaction, Proceedings of the National Academy of Sciences 116, 26523–26533.

(42.) Stefan, N., Martin-Killias, P., Wyss-Stoeckle, S., Honegger, A., Zangemeister-Wittke, U., and Plückthun, A. (2011) DARPins Recognizing the Tumor-Associated Antigen EpCAM Selected by Phage and Ribosome Display and Engineered for Multivalency, Journal of Molecular Biology 413, 826–843.

(43.) Gottschalk, N., Kimmig, R., Lang, S., Singh, M., and Brandau, S. (2012) Anti-Epidermal Growth Factor Receptor (EGFR) Antibodies Overcome Resistance of Ovarian Cancer Cells to Targeted Therapy and Natural Cytotoxicity, In International Journal of Molecular Sciences, pp 12000–12016.

(44.) Sewell, J. M., Macleod, K. G., Ritchie, A., Smyth, J. F., and Langdon, S. P. (2002) Targeting the EGF receptor in ovarian cancer with the tyrosine kinase inhibitor ZD 1839 (‘Iressa’), British Journal of Cancer 86, 456–462.

(45.) Anil-Inevi, M., Sağlam-Metiner, P., Kabak, E. C., and Gulce-Iz, S. (2020) Development and verification of a three-dimensional (3D) breast cancer tumor model composed of circulating tumor cell (CTC) subsets, Molecular Biology Reports 47, 97–109.

(46.) Prang, N., Preithner, S., Brischwein, K., Göster, P., Wöppel, A., Müller, J., Steiger, C., Peters, M., Baeuerle, P. A., and da Silva, A. J. (2005) Cellular and complement-dependent cytotoxicity of Ep-CAM-specific monoclonal antibody MT201 against breast cancer cell lines, British Journal of Cancer 92, 342–349.

(47.) Yang, J., Isaji, T., Zhang, G., Qi, F., Duan, C., Fukuda, T., and Gu, J. (2020) EpCAM associates with integrin and regulates cell adhesion in cancer cells, Biochemical and Biophysical Research Communications 522, 903–909.

(48.) Aigner, A., Hsieh, S. S., Malerczyk, C., and Czubayko, F. (2000) Reversal of HER-2 over-expression renders human ovarian cancer cells highly resistant to taxol, Toxicology 144, 221–228.

(49.) Wang, W., Gao, Y., Hai, J., Yang, J., and Duan, S. (2019) HER2 decreases drug sensitivity of ovarian cancer cells via inducing stem cell-like property in an NFκB-dependent way, Bioscience Reports 39, BSR20180829.

(50.) Conlon, N. T., Kooijman, J. J., van Gerwen, S. J. C., Mulder, W. R., Zaman, G. J. R., Diala, I., Eli, L. D., Lalani, A. S., Crown, J., and Collins, D. M. (2021) Comparative analysis of drug response and gene profiling of HER2-targeted tyrosine kinase inhibitors, British Journal of Cancer 124, 1249–1259.

(51.) Stanley, A., Ashrafi, G. H., Seddon, A. M., and Modjtahedi, H. (2017) Synergistic effects of various Her inhibitors in combination with IGF-1R, C-MET and Src targeting agents in breast cancer cell lines, Scientific Reports 7, 3964.

(52.) Smith, S. E., Mellor, P., Ward, A. K., Kendall, S., McDonald, M., Vizeacoumar, F. S., Vizeacoumar, F. J., Napper, S., and Anderson, D. H. (2017) Molecular characterization of breast cancer cell lines through multiple omic approaches, Breast Cancer Research 19, 65.

(53.) Vranic, S., Gatalica, Z., and Wang, Z.-Y. (2011) Update on the molecular profile of the MDA-MB-453 cell line as a model for apocrine breast carcinoma studies, Oncol Lett 2, 1131–1137.

(54.) Idibie, C. A., Davids, H., and Iyuke, S. E. (2007) Cytotoxicity of purified cassava linamarin to a selected cancer cell lines, Bioprocess and Biosystems Engineering 30, 261–269.

(55.) Kousparou, C. A., Epenetos, A. A., and Deonarain, M. P. (2002) Antibody-guided enzyme therapy of cancer producing cyanide results in necrosis of targeted cells, International Journal of Cancer 99, 138–148.

(56.) Li, J., Li, H., Zhu, L., Song, W., Li, R., Wang, D., and Dou, K. (2010) The adenovirus-mediated linamarase/linamarin suicide system: A potential strategy for the treatment of hepatocellular carcinoma, Cancer Letters 289, 217–227.

(57.) Mkpong, O. E., Yan, H., Chism, G., and Sayre, R. T. (1990) Purification, Characterization, and Localization of Linamarase in Cassava 1, Plant Physiology 93, 176–181.

(58.) Bhattacharya, R., and Lakshmana Rao, P. V. (1997) Cyanide induced DNA fragmentation in mammalian cell cultures, Toxicology 123, 207–215.

(59.) Marziaz, M. L., Frazier, K., Guidry, P. B., Ruiz, R. A., Petrikovics, I., and Haines, D. C. (2013) Comparison of brain mitochondrial cytochrome c oxidase activity with cyanide LD50 yields insight into the efficacy of prophylactics, Journal of Applied Toxicology 33, 50–55.

(60.) Borowitz, J. L., Rathinavelu, A., Kanthasamy, A., Wilsbacher, J., and Isom, G. E. (1994) Accumulation of Labeled Cyanide in Neuronal Tissue, Toxicology and Applied Pharmacology 129, 80–85.

(61.) Luisa Cortés, M., García-Escudero, V., Hughes, M., and Izquierdo, M. (2002) Cyanide bystander effect of the linamarase/linamarin killer-suicide gene therapy system, The Journal of Gene Medicine 4, 407–414.

(62.) Colella, P., Ronzitti, G., and Mingozzi, F. (2018) Emerging Issues in AAV-Mediated In Vivo Gene Therapy, Molecular Therapy - Methods & Clinical Development 8, 87–104.

(63.) Shi, W., and Bartlett, J. S. (2003) RGD inclusion in VP3 provides adeno-associated virus type 2 (AAV2)-based vectors with a heparan sulfate-independent cell entry mechanism, Molecular Therapy 7, 515–525.

(64.) Theuerkauf, S. A., Herrera-Carrillo, E., John, F., Zinser, L. J., Molina, M. A., Riechert, V., Thalheimer, F. B., Börner, K., Grimm, D., Chlanda, P., Berkhout, B., and Buchholz, C. J. (2023) AAV vectors displaying bispecific DARPins enable dual-control targeted gene delivery, Biomaterials 303, 122399.

(65.) Tan, T. K., Rijal, P., Rahikainen, R., Keeble, A. H., Schimanski, L., Hussain, S., Harvey, R., Hayes, J. W. P., Edwards, J. C., McLean, R. K., Martini, V., Pedrera, M., Thakur, N., Conceicao, C., Dietrich, I., Shelton, H., Ludi, A., Wilsden, G., Browning, C., Zagrajek, A. K., Bialy, D., Bhat, S., Stevenson-Leggett, P., Hollinghurst, P., Tully, M., Moffat, K., Chiu, C., Waters, R., Gray, A., Azhar, M., Mioulet, V., Newman, J., Asfor, A. S., Burman, A., Crossley, S., Hammond, J. A., Tchilian, E., Charleston, B., Bailey, D., Tuthill, T. J., Graham, S. P., Duyvesteyn, H. M. E., Malinauskas, T., Huo, J., Tree, J. A., Buttigieg, K. R., Owens, R. J., Carroll, M. W., Daniels, R. S., McCauley, J. W., Stuart, D. I., Huang, K.-Y. A., Howarth, M., and Townsend, A. R. (2021) A COVID-19 vaccine candidate using SpyCatcher multimerization of the SARS-CoV-2 spike protein receptor-binding domain induces potent neutralising antibody responses, Nature Communications 12, 542.

(66.) Gibson, D. G., Young, L., Chuang, R.-Y., Venter, J. C., Hutchison, C. A., and Smith, H. O. (2009) Enzymatic assembly of DNA molecules up to several hundred kilobases, Nature Methods 6, 343–345.

(67.) Xiao, X., Li, J., and Samulski Richard, J. (1998) Production of High-Titer Recombinant Adeno-Associated Virus Vectors in the Absence of Helper Adenovirus, Journal of Virology 72, 2224–2232.

