## Supplementary Information for "Genetically encoded SpyTag enables modular AAV retargeting via SpyCatcher-fused ligands for targeted gene delivery"

[4] Current address: Prolific Machines, Emeryville, CA 94608, United States

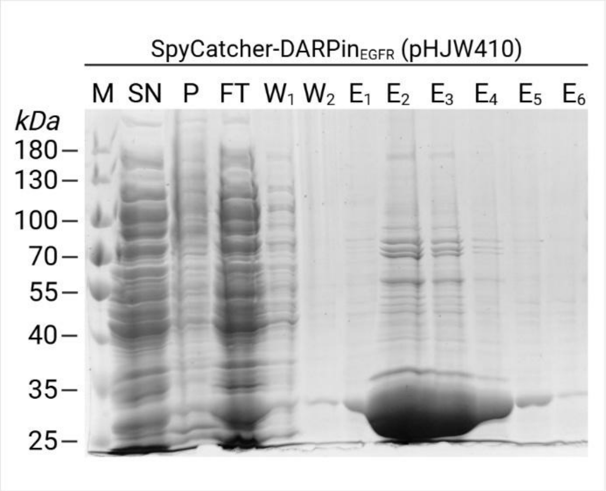


**Figure S1: Analysis of SpyC-DARPin$_{EGFR}$ purification by SDS-PAGE and Coomassie staining.**

SpyCatcher-DARPin$_{EGFR}$ was produced in *E. coli* from plasmid pHJW410 and purified by IMAC. Samples from different purification steps were subjected to analysis by SDS-PAGE followed by Coomassie staining. M, protein size marker; SN, soluble fraction of lysate; P, insoluble fraction of lysate; FT, flow through; W, wash; E, elution after purification. Calculated SpyCatcher-DARPin$_{EGFR}$ mass: 27.0 kDa.

```
SpyCatcher001    1     MAGVDTLSGLSSEQGQSGDMTIEEDSATHIKFSKRDEDGKELAGATMELRDSSGKTISTWISDGQVKDFY
SpyCatcher001Δ   1     ---------------------MSGDSATHIKFSKRDEDGKELAGATMELRDSSGKTISTWISDGQVKDFY
SpyCatcher003    1     --MVTTLSGLSGEQGPSGDMTTEEDSATHIKFSKRDEDGRELAGATMELRDSSGKTISTWISDGHVKDFY
SpyCatcher003Δ   1     ---------------------MSGDSATHIKFSKRDEDGRELAGATMELRDSSGKTISTWISDGHVKDFY

SpyCatcher001    71    LYPGKYTFVETAAPDGYEVATAITFTVNEQGQVTVNGKATKGDAHI
SpyCatcher001Δ   50    LYPGKYTFVETAAPDGYEVATAITFTVNEQGQVTVNG---------
SpyCatcher003    69    LYPGKYTFVETAAPDGYEVATPIEFTVNEDGQVTVDGEATEGDAHT
SpyCatcher003Δ   50    LYPGKYTFVETAAPDGYEVATPIEFTVNEDGQVTVDG---------
```

**Figure S2: Generation of SpyCatcher003Δ by sequence comparison with SpyCatcher001Δ.**

SpyCatcher003Δ was generated after comparing the protein sequences of SpyCatcher001 and SpyCatcher001Δ, identifying the truncated regions in SpyCatcher001Δ and applying the same truncations to the parental SpyCatcher003 sequence to finally yield SpyCatcher003Δ. Consensus sequence is marked blue. Alignment was generated using MUSCLE in SnapGene.

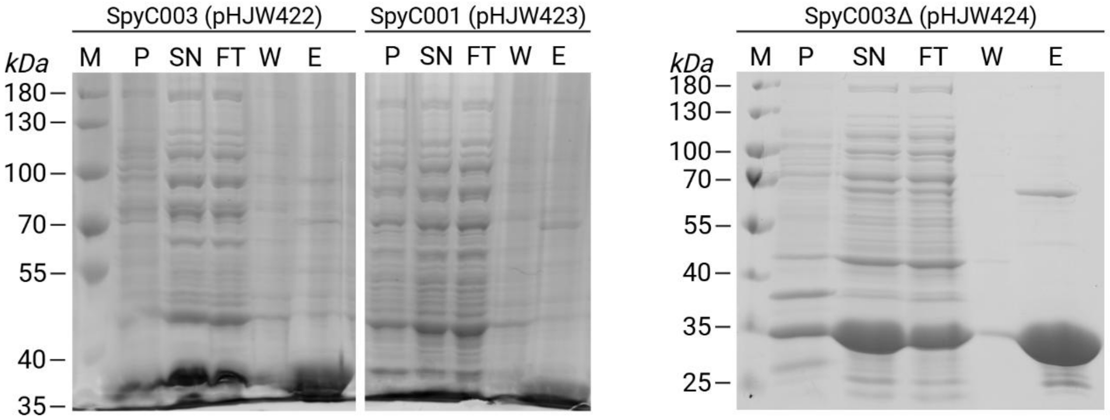


**Figure S3: Analysis of SpyCatcher variants purification by SDS-PAGE and Coomassie staining.**

SpyCatcher003-DARPin$_{EGFR}$, SpyCatcher001-DARPin$_{EGFR}$, SpyCatcher003Δ-DARPin$_{EGFR}$ were produced in *E. coli* from plasmids pHJW422, pHJW423 and pHJW424, respectively and purified by IMAC. Samples from different purification steps were subjected to analysis by SDS-PAGE followed by Coomassie staining. M, protein size marker; SN, soluble fraction of lysate; P, insoluble fraction of lysate; FT, flow through; W, wash; E, elution after purification. Calculated masses: SpyCatcher003-DARPin$_{EGFR}$, 30.2 kDa; SpyCatcher001-DARPin$_{EGFR}$, 30.5 kDa; SpyCatcher001-DARPin$_{EGFR}$, 27.3 kDa.

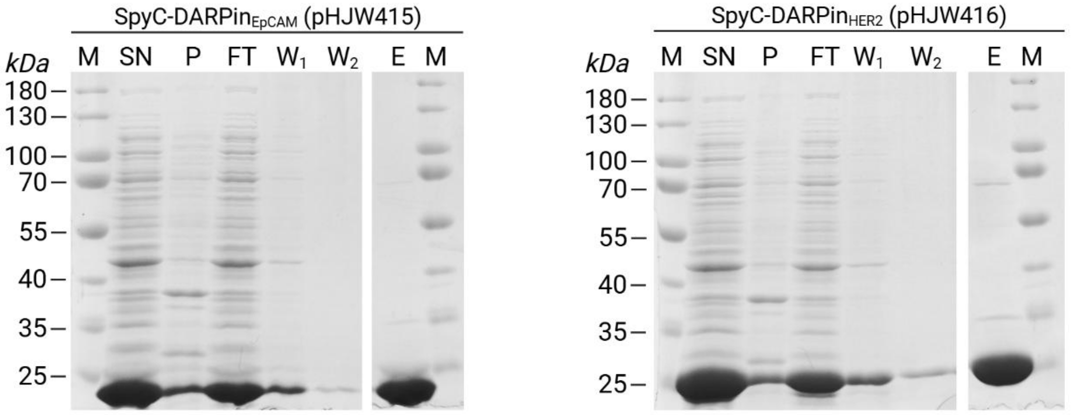


**Figure S4: Analysis of SpyCatcher-DARPin$_{EpCAM}$ and SpyCatcher-DARPin$_{HER2}$ purification by SDS-PAGE and Coomassie staining.**

SpyCatcher-DARPin$_{EpCAM}$ and SpyCatcher-DARPin$_{HER2}$ were produced in *E. coli* from plasmids pHJW415 and pHJW416, respectively and purified by IMAC. Samples from different purification steps were subjected to analysis by SDS-PAGE followed by Coomassie staining. M, protein size marker; SN, soluble fraction of lysate; P, insoluble fraction of lysate; FT, flow through; W, wash; E, elution after purification. Calculated masses: SpyCatcher-DARPin$_{EpCAM}$ , 27.5 kDa; SpyCatcher-DARPin$_{HER2}$ , 27.4 kDa.

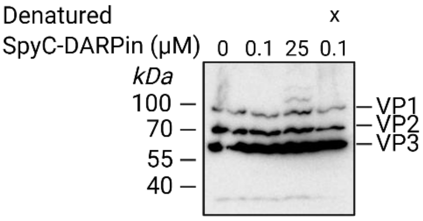


**Figure S5: Characterization of LIN-mScarlet-AAV**

Western blot analysis of PEG-precipitated viral capsid proteins VP1, VP2 and VP3 of SpyT453-AAV carrying linamarase-mScarlet from cell culture supernatant. Numbers indicate the concentration of SpyCatcher-DARPin$_{EGFR}$ protein during incubation. “x” denotes AAV denaturation by boiling at 98 °C for 10 min before SC-DARPin coupling. SpyT-AAV and SpyC-DARPin$_{EGFR}$ concentrations are listed in Table S2.

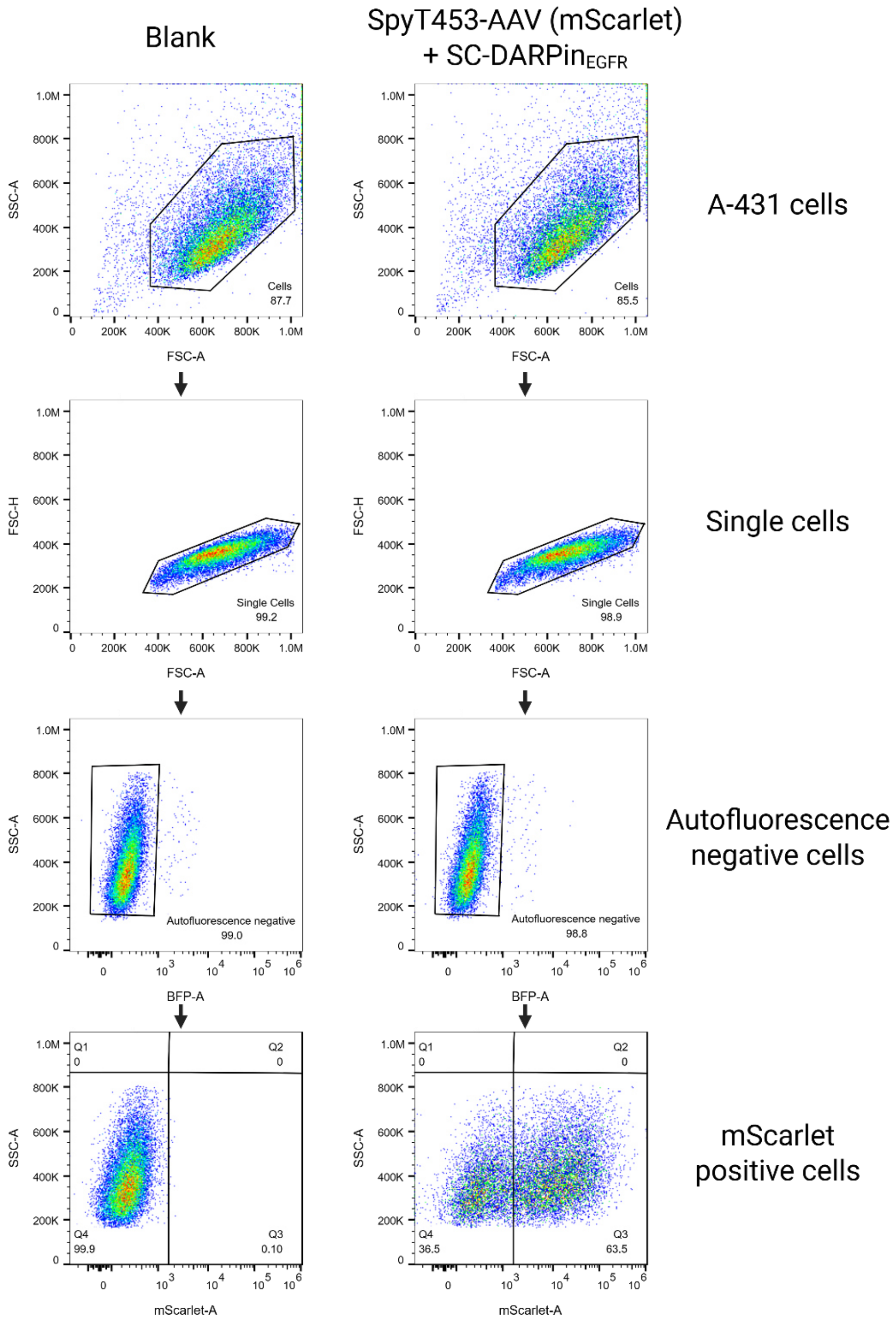


**Figure S6: Gating strategy of AAV transduction experiments analyzed by flow cytometry.**

The gating strategy of representative samples from experiments performed with SpyTag-AAVs on various cancer cell lines (depicted here: A-431 cells) is shown. First, cells were gated based on FSC-A and SSC-A signal, then doublets were excluded based on FSC-H and FSC-A signal. Autofluorescent cells were excluded based on BFP-A signal and mScarlet positive cells were gated based on untransduced cells (Blank). Data corresponds to Fig 3 in the main text.

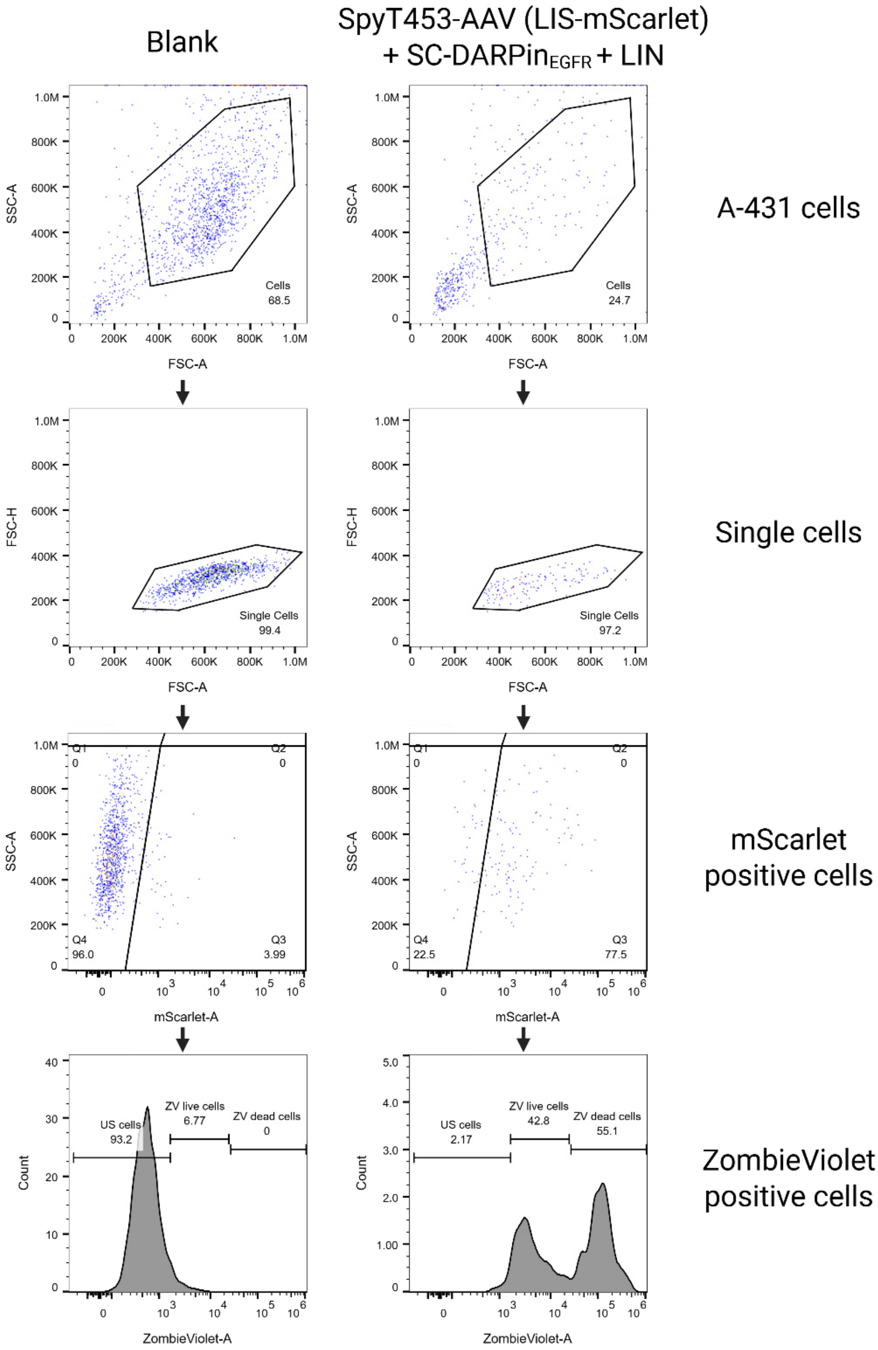


**Figure S7: Gating strategy of AAV transduction experiments with live/dead staining analyzed by flow cytometry.**

The gating strategy of representative samples from experiments performed with SpyTag-AAVs and live/dead staining of A-431 cells is shown. Cells were gated based on FSC-A and SSC-A signal, before excluding doublets based on FSC-H and FSC-A signal. mScarlet positive cells were gated based on untransduced samples (Blank). Zombie Violet positive cells were gated based on unstained samples (Blank). Data corresponds to Fig. 6 In the main text. Abbreviations: LIN, Linamarin; LIS, Linamarase.

**Table S1: SpyT-AAV and SpyC-DARPin$_{EGFR}$ concentrations in the Western blot.**

Concentrations of SpyT-AAV and SpyC-DARPin$_{EGFR}$ during coupling (sample volume: 40 µl) and after loading onto SDS-PAGE (volume: 10 µl). Data corresponds to Figure 2 in the main text.

| | Coupling with SpyC-DARPin$_{EGFR}$ | | SDS-PAGE and Western blot | |
|---|---|---|---|---|
| | AAV (vg/ml) | SpyC-DARPin (µM) | AAV (vg) | SpyC-DARPin (nmol) |
| SpyT587-AAV | $2.5*10^{10}$ | 42.0 | $2.0*10^{8}$ | 0.336 |
| SpyT453-AAV | $1.1*10^{10}$ | 42.0 | $9.0*10^{7}$ | 0.336 |
| SpyT587VP2-AAV | $5.3*10^{10}$ | 42.0 | $4.3*10^{8}$ | 0.336 |
| SpyT453VP2-AAV | $7.3*10^{10}$ | 42.0 | $5.8*10^{8}$ | 0.336 |

**Table S2: SpyT-AAV and SpyC-DARPin$_{EGFR}$ concentrations in the Western blot.**
Concentrations of SpyT453-AAV (carrying linamarase-mScarlet) and SpyC-DARPin$_{EGFR}$ during coupling (sample volume: 20 µl) and after loading onto SDS-PAGE (volume: 10 µl). Data corresponds to Figure S5 in the supplementary information.

| | Coupling with SpyC-DARPin$_{EGFR}$ | | SDS-PAGE and Western blot | |
|---|---|---|---|---|
| | AAV (vg/ml) | SpyC-DARPin (µM) | AAV (vg) | SpyC-DARPin (nmol) |
| Lane 1 | $5.5*10^{9}$ | - | $4.4*10^{7}$ | - |
| Lane 2 | $5.5*10^{9}$ | 0.1 | $4.4*10^{7}$ | 0.000807 |
| Lane 3 | $5.5*10^{9}$ | 25.2 | $4.4*10^{7}$ | 0.202 |
| Lane 4 | $5.5*10^{9}$ | 0.1 | $4.4*10^{7}$ | 0.000807 |

**Table S3: Conversion efficiency of Linamarin into HCN.**
Conversion efficiency of Linamarin into HCN was calculated from mean quantified HCN in linamarase transduced and linamarin treated samples. Data corresponds to Figure 6 in the main text.

| Linamarin (µg/ml) | Linamarin (µmol/ml) | Mean HCN (µg/ml) | Mean HCN (µmol/ml) | Conversion Efficiency (%) |
|---|---|---|---|---|
| 0 | 0.0000 | 0.2779 | 0.0103 | |
| 250 | 1.0112 | 2.5846 | 0.0956 | 9.46 |
| 500 | 2.0224 | 5.3071 | 0.1963 | 9.71 |
| 750 | 3.0336 | 7.7827 | 0.2879 | 9.49 |
| 1000 | 4.0448 | 10.8946 | 0.4031 | 9.96 |
| 1500 | 6.0672 | 15.3020 | 0.5661 | 9.33 |
| 2000 | 8.0896 | 24.0579 | 0.8900 | 11.00 |
| | | | Mean (%) | 9.83 |
| | | | SD (%) | 0.62 |

**Table S4: Plasmids used and generated in this study.**

| Category | Plasmid Name | Description | Backbone/ Reference |
|---|---|---|---|
| AAV plasmids | AdH | Promoter-E2A-E4-VA-RNA | pHelper vector, Cell Biolabs (Cat.No. VPK-402, Part No. 340202) |
| | pMH321 | pRC2-587(SpyTag)-R585/588A | pAAV-RC2, Cell Biolabs (Cat.No. VPK-402, Part No. VPK-422) |
| | pHJW414 | pRC2-453(SpyTag)-R585/588A | pAAV-RC2, Cell Biolabs (Cat.No. VPK-402, Part No. VPK-422) |
| | pHJW162 | pR2-VP1/3(R585/588A) | pVP1/3 *(1)* |
| | pHJW341 | pCMV-VP2-587(SpyTag)-R585/588A-VP3KO | pEGFP-C3 (Clontech) |
| | pHJW351 | pCMV-VP2-453(SpyTag)-R585/588A-VP3KO | pEGFP-C3 (Clontech) |
| | CMV-mScarlet | ITR-pCMV-mScarlet-ITR | pCMV-mScarlet (D. Grimm) |
| | pHJW427 | ITR-pCMV-Linamarase-2A-mScarlet-ITR | pCMVmScarlet (D. Grimm), Linamarase: pWW315 *(2)* |
| SpyCatcher plasmids | pHJW410 | pT7-SpyCatcher001Δ-GSS-Linker-DARPin_E01-$His_6$ | pRSET, SpyCatcher-Toolbox1664, pHJW156 *(3)* |
| | pHJW415 | pT7-SpyCatcher001Δ-GSS-Linker-DARPin_Ec1-$His_6$ | pRSET, SpyCatcher-Toolbox1664, pMH327 *(3)* |
| | pHJW416 | pT7-SpyCatcher001Δ-GSS-Linker-DARPin_9.29-$His_6$ | pRSET, SpyCatcher-Toolbox1664, pMH328 *(3)* |
| | pHJW422 | pT7-SpyCatcher003-GSS-Linker-DARPin_E01-$His_6$ | pRSET, pOSY115 *(3)* |
| | pHJW423 | pT7-SpyCatcher001-GSS-Linker-DARPin_E01-$His_6$ | pRSET, SpyCatcher-Toolbox1664 |
| | pHJW424 | pT7-SpyCatcher003Δ-GSS-Linker-DARPin_E01-$His_6$ | pRSET, pOSY115 *(3)* |

**Table S5: AAV composition.**

| Name | Description | Plasmids |
|---|---|---|
| SpyT587-AAV (mScarlet) | pRC2(R585/588A)(587-SpyTag) AAV | AdH<br>pMH321<br>CMV-mScarlet |
| SpyT453-AAV (mScarlet) | pRC2(R585/588A)(453-SpyTag) AAV | AdH<br>pHJW414<br>CMV-mScarlet |
| SpyT587VP2-AAV (mScarlet) | pR2-VP1/3(R585/588A) + VP2(587-SpyTag) | AdH<br>pHJW162<br>pHJW341<br>CMV-mScarlet |
| SpyT453VP2-AAV (mScarlet) | pR2-VP1/3(R585/588A) + VP2(587-SpyTag) | AdH<br>pHJW162<br>pHJW351<br>CMV-mScarlet |
| SpyT453-AAV (linamarase-mScarlet) | pRC2(R585/588A)(453-SpyTag) AAV | AdH<br>pHJW414<br>pHJW427 |

**References**


(1.) Gomez, E. J., Gerhardt, K., Judd, J., Tabor, J. J., and Suh, J. (**2016**) Light-Activated Nuclear Translocation of Adeno-Associated Virus Nanoparticles Using Phytochrome B for Enhanced, Tunable, and Spatially Programmable Gene Delivery, *ACS Nano 10*, 225-237.

(2.) Link, N., Aubel, C., Kelm, J. M., Marty, R. R., Greber, D., Djonov, V., Bourhis, J., Weber, W., and Fussenegger, M. (**2006**) Therapeutic protein transduction of mammalian cells and mice by nucleic acid-free lentiviral nanoparticles, *Nucleic Acids Research 34*, e16-e16.

(3.) Hörner, M., Jerez-Longres, C., Hudek, A., Hook, S., Yousefi, O. S., Schamel, W. W. A., Hörner, C., Zurbriggen, M. D., Ye, H., Wagner, H. J., and Weber, W. (**2021**) Spatiotemporally confined red light-controlled gene delivery at single-cell resolution using adeno-associated viral vectors, *Science Advances 7*, eabf0797.
